# Genes mirror migrations and cultures in prehistoric Europe – a population genomic perspective

**DOI:** 10.1101/072926

**Authors:** Torsten Günther, Mattias Jakobsson

## Abstract

Genomic information from ancient human remains is beginning to show its full potential for learning about human prehistory. We review the last few years' dramatic finds about European prehistory based on genomic data from humans that lived many millennia ago and relate it to modern-day patterns of genomic variation. The early times, the Upper Palaeolithic, appears to contain several population turn-overs followed by more stable populations after the Last Glacial Maximum and during the Mesolithic. Some 11,000 years ago the migrations driving the Neolithic transition start from around Anatolia and reach the north and the west of Europe millennia later followed by major migrations during the Bronze age. These findings show that culture and lifestyle were major determinants of genomic differentiation and similarity in pre-historic Europe rather than geography as is the case today.

---

The genetic makeup of modern-day European groups and the historical processes which generated these patterns of population structure and isolation-by-distance have attracted great interest from population geneticists, anthropologists, historians and archaeologists. Early studies of genetic variation among Europeans indicated that the major axes of variation are highly correlated with geography [1], a pattern that has been confirmed and refined during the age of population-genomics (Figure 1A) [2, 3]. Today, we have learned from studying the genomics of past peoples of Europe that the modern-day patterns of genomic variation were shaped by several major demographic events in the past. These events include the first peopling of Europe [4, 5], the Neolithic transition [6], and later migrations during the Bronze Age [7, 8].

**Figure 1:**
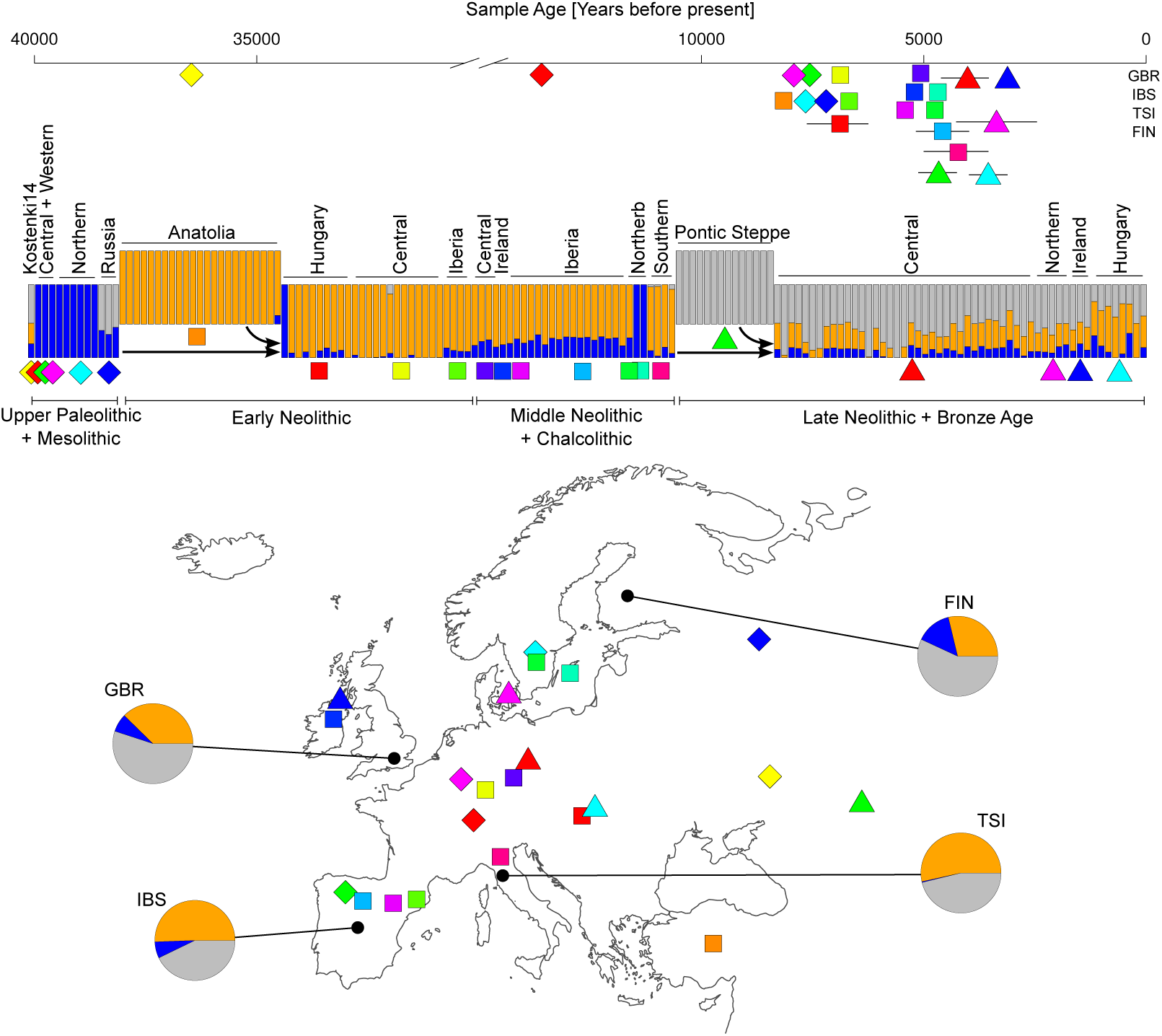
Principal component analyses for different time periods and focusing on western Eurasia. (A) Modern-day samples, (B) Upper Palaeolithic and Mesolithic samples, including samples identified as representative for prehistoric clusters in [5], arrows show transitions during the Upper Palaeolithic, (C) early and middle Neolithic samples, and (D) late Neolithic and Bronze Age samples. Individuals from previous sub-figures are depicted
in gray.

Recent years have seen technological and methodological advances which gave rise to the field of ancient genomics. By studying the genomes of individuals from the past, we are able to investigate individuals (and groups) that lived during those times and during particular events and periods. This new approach has provided unprecedented power and resolution to study human prehistory. In contrast to prior views of population continuity, we now know that migrations (from various areas) were numerous during the past and that they shaped modern-day patterns of variation, in combination with isolation-by-distance processes. European prehistory has proven to be more complex than what would be concluded from the most parsimonious models based on modern-day population-genomic data [9]. Here, we review the most recent findings about European prehistory based on ancient genomics.

## 1. European Hunter-Gatherers during the Upper Palaeolithic and the Mesolithic

The first anatomically modern humans spread over Europe around 45,000 years ago [10, 11, 12, 13] where they met [11] and mixed with Neandertals [14, 15, 16]. The contributions of these early Europeans to modern populations appear limited [16, 5], and mitochondrial data shows that some haplogroups found in some individuals who lived in Europe during the Upper Palaeolithic are not present in modern-day Europeans, which suggests loss of some genetic variation or replacement through later migrations [5, 17]. However, European individuals that lived after 37,000 years ago, in the Upper Palaeolithic, have been suggested to already carry major genetic components found in today’s Europe [4, 5]. The peoples living in Europe during these times were obviously influenced by climatic changes, at least in terms of which geographic areas that were habitable. For instance, during the Last Glacial Maximum (LGM) [26,500 to 19,000 years ago, 18], the north of Europe became uninhabitable and the remaining habitable areas became fragmented [19, 20]. The LGM likely caused a severe bottleneck for the groups at the time [21], when people may have moved south into specific refugia, which is indicated by low genomic diversity of later hunter-gatherer groups [22] and reduced mitochondrial diversity [17]. After the LGM, migrants from southeast appear in Europe and mixed with the local European post ice age groups [5] resulting in a population that may have remained largely uniform during the late Upper Paleolithic and possibly into the Mesolithic in some areas [23, 24, 25, 5], although this interpretation is still uncertain as only a limited number of geographic areas in Europe have been investigated. The genomic make-up of these hunter-gatherers fall outside of the genetic variation of modern-day people, including western Eurasians (Figure 1B). The genomic patterns of western [23, 26] and central European Mesolithic individuals [24, 25] appear distinct from eastern European Mesolithic individuals who showed a stronger influence from ancient north Eurasian populations [7, 27, 5] (Figure 2). Scandinavian Mesolithic individuals [22, 24] show genetic affinities to both the western and the eastern Mesolithic groups, suggesting a continuum of genetic variation, possibly caused by admixture from both geographic areas. Such a scenario would be consistent with patterns seen in many animal and plant species which recolonized the Scandinavian peninsula from the south and north-east after the last glacial period [28, 29, 30]. Surprisingly, some of these Scandinavian hunter-gatherers were carriers of the derived allele in the EDAR gene [27]. EDAR affects hair thickness and teeth morphology and has been subject to a selective sweep in East Asia, but the derived allele is virtually absent from modern day Europe [27]. Together with evidence from stone tool technology [31] and the potential introduction of domesticated dogs from Asia [32, 33, 34] this suggests large scale networks across Eurasia already during the Upper Palaeolithic and Mesolithic periods.

**Figure 2:**
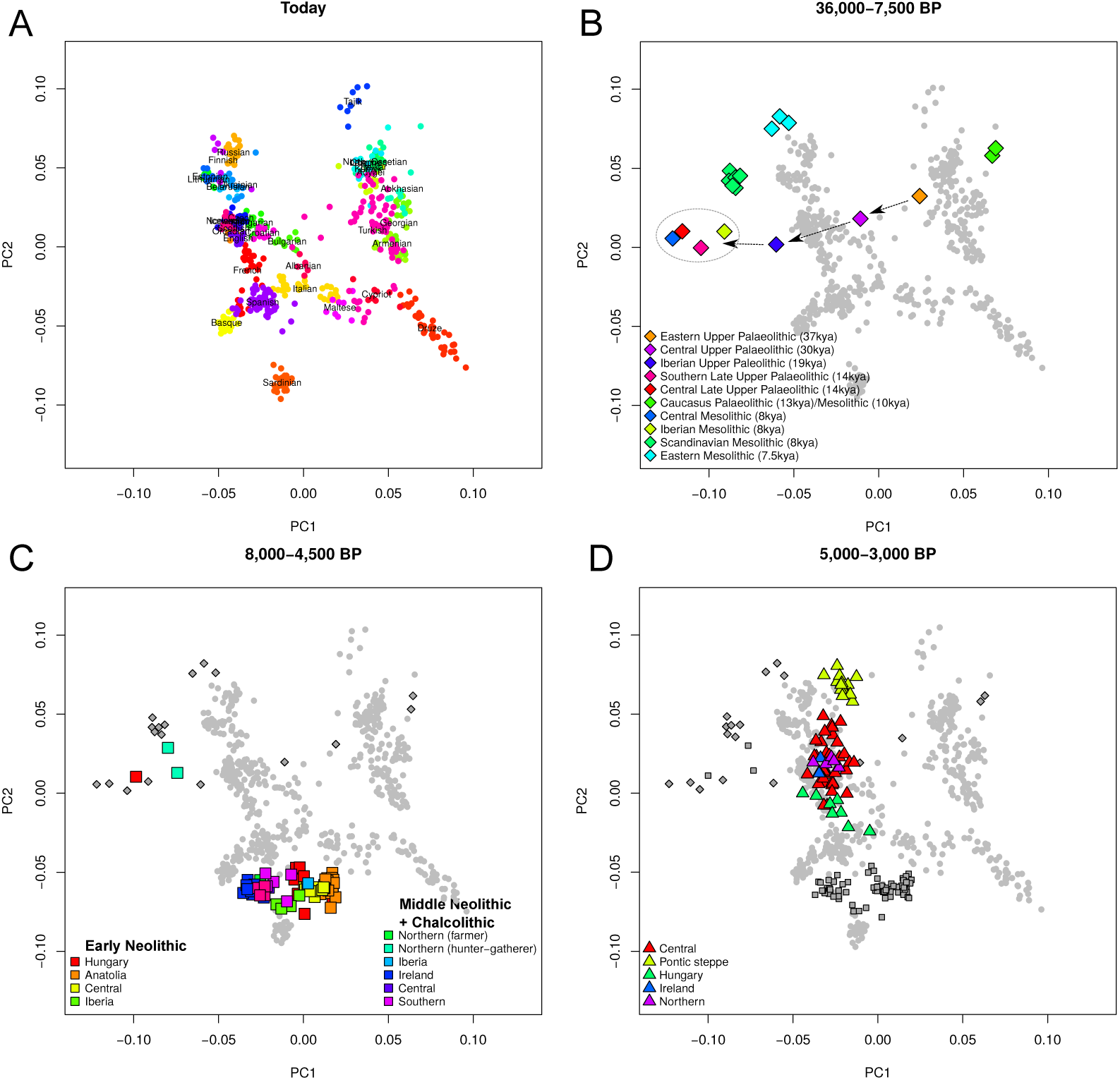
Supervised Admixture [75] analysis of ancient samples and modern populations from the 1000 genomes project [76] shown together with the approximate geographic origin of the ancient samples and the sample age. Admixture was run in supervised mode using Western hunter-gatherers, Anatolian Neolithic farmers and Yamnaya herders from the Pontic steppe as “source” populations [7]. Note that these three “source” groups were themselves mixed groups (see text). Pie charts show admixture proportions for modern European populations from the 1000 genomes project.

## 2. The Neolithic transition

The Neolithic transition – the transformation from a hunter-gatherer lifestyle to a sedentary farming lifestyle – has been an exceptionally important change in human history and forms the basis for emergence of civilizations. This transition occurred independently in different parts of the world [35]. For western Eurasia, the first evidence of farming practices have been found in the fertile crescent and dated to 11,000-12,000 years BP. From there, farming spread into Anatolia and Europe, reaching Scandinavia and the British Isles around 6,000 years ago. It has been a long-standing debate whether farming practices spread as an idea [“cultural diffusion”, 36, 37] or via migration of people [“demic diffusion”, 38]. Genetic studies using uniparental markers could be interpreted as supporting both theories [e.g. 39, 40, 41, 42, 43, 44, 45, 46, 47, 48, 49], but genomic data from early farmers from different parts of Europe clearly showed a strong differentiation between them and European hunter-gatherers [6, 22, 24, 50, 51, 52, 53]. The differences among the two groups were almost as strong as between modern-day populations from different continents [22]. The genetic composition of the Mesolithic individuals falls outside the range of modern-day Eurasians, but they are genetically most similar to modern populations from north and northeast of Europe. Early farmers exhibit marked genetic similarities with modern-day southwestern Europeans, but not with modern-day groups from the Near and Middle East (Figure 1C) [6, 22, 24]. The affinity of Neolithic Europeans to modern Southern Europeans is particularly pronounced for the population isolates of Sardinia (see Box 1) and Basques [52]. The connection to Basques is surprising since they were long considered to be more-or-less direct descendants of Mesolithic Europeans [54, 55] (see Box 1).

### Box 1 European population “outliers” and farming out of Sardinia?

The Basque people have been suggested to be descendants of Paleolithic and Mesolithic groups based on language and early genetic studies [72, 54, 73, 55]. However, also the Basque have been shown to be descendants of early European farmers who mixed with resident hunter-gatherers [52]. The reason why the Basque stood out genetically is the lack of gene-flow in the last couple of millennia when most European groups have been affected by several migrations during the Bronze Age and historic times [7, 8]. Basques remained relatively isolated during these times.

Some four decades ago, Luca Cavalli-Sforza and colleagues pioneered the use of genetic data to investigate European demographic history [1, 54]. For instance, they challenged the dominant theory that farming practices spread as an idea during the Neolithic transition and instead suggested that farming groups replaced local hunter-gatherers [1, 54]. Although the data was limited at the time, and that the patters observed were also consistent with other scenarios that do not involve mass-migration [68], the work of Cavalli-Sforza and colleagues pioneered today’s powerful approaches of using genomic variation to infer human history. Cavalli-Sforza and colleagues highlighted some “outlier” populations as being of particular interest: Sardinians, Basques, and Saami. Interestingly, the processes that led to the specific genetic patterns in these three groups differ markedly. When the first genomic data from early European farmers was obtained from a Scandinavian skeleton and a mummy from the Alps – commonly known as Ötzi or the Tyrolean Iceman – researchers were surprised that they showed a strong population affinity to modern-day Sardinians [6, 69]. This observation was even more puzzling since archaeo-logical investigations demonstrate that farming started in the Fertile Crescent [59]. Later studies confirmed those affinities for early farmers from the areas of modern-day Sweden [22], Germany [24], Hungary [50], Spain [52, 51] and Ireland [53], which all showed particularly strong affinities to modern-day Sardinians and not to modern-day Near Eastern populations. Ancient genomic data from Neolithic Anatolia [27, 56, 58] resolved the puzzle by showing that Neolithic individuals from Anatolia were genetically similar to Neolithic farmers from across Europe and modern-day Sardinians. A likely scenario would be: i) The group of early farmers who migrated westwards and northward settled on Sardinia where they encountered (and admixed with) few or no hunter-gatherers; ii) Sardinians remained relatively isolated from migrations that largely affected the rest of Europe [70]. The Northern European Saami are not particularly related to the first pioneers that colonized northern Europe after the LGM, nor the Neolithic groups arriving a few millennia later [22] as suggested by some archaeological studies [71]. Instead they appear to be related to groups that migrated from the east to northern Europe in more recent times, although much is still unknown about the Saami as no genetic data has been generated from pre-historic human remains of the very north of Scandinavia – Sápmi.

Later studies on early farmers from Anatolia and the Levant identified them as source population of the European Neolithic groups [27, 56, 57, 58], which is well in-line with long standing archaeological results [59]. The first farmers of central Anatolia lived in small transition groups 10,000 years BP with early pre-pottery Neolithic practices including small-scale cultivation, but heavily relying on foraging as well [58]. Such groups formed the genetic basis of the farmers that expanded first within Anatolia and the Near East and then started expanding into Europe [58, 57] in contrast to groups in e.g. the eastern Fertile Crescent [60, 57, 61] that contributed limited genetic material to the early European farmers. Demographic changes since that period in Anatolia, such as gene-flow from the east, explain the discrepancy between modern-day Anatolian genetic make-up and Neolithic groups of the area [56].

Archaeological investigations have also suggested that farming spread through two different routes across the European continent: one route along the Danube river into central Europe and one along the Mediterranean coast [62]. Ancient DNA from different parts of Europe is also consistent with such a dichotomy [63, 64, 53].

Farming practices probably allowed to maintain larger groups of people, which is indicated by a substantially higher genetic diversity of farming groups [22, 50]. Hunter-gatherers and farmers, however, did not remain isolated from each other. The two groups must have met and mixed, which can be seen in middle Neolithic farmers that display distinct and increased fractions of genetic ancestry from hunter-gatherer groups [22, 7, 52]. The finding that hunter-gatherers and farmers mixed shows that neither the strict demic nor the strict cultural diffusion model accurately describe the events during the Neolithic transition, a model that some archaeologists had previously favored [65, 66]. The hunter-gatherer lifestyle was eventually replaced by farming, but the farming groups assimilated hunter-gatherers resulting in differently admixed groups across Europe, a pattern that can still be seen in current-day Europeans [22, 24] (Figure 2). An interesting case-example is the KO1 individual recovered from an archaeological site with a clear farming context in modern-day Hungary, who was genetically very similar to European hunter-gatherers [50]. This KO1 individual is possibly a first generation (hunter-gatherer) migrant into the farming community. More generally, farmers from the middle Neolithic and early Chalcolithic periods (6,000 to 4,500 years ago) display additional admixture from Mesolithic hunter-gatherers compared to early Neolithic groups [7, 52] (Figure 2). The positive correlation of hunter-gatherer related admixture in farming groups with time [52] implies that people of both groups have met and mixed at different points in time and in different parts of Europe. The process of admixture must have continued for at least two millennia which raises the question where the genetically hunter-gatherer populations were living during the Neolithic. Some scholars suggest that those groups retreated to the Atlantic coast during the early Neolithic from where they re-surged later on [e.g. 49, 7]. It has also been suggested that some hunter-gatherer groups still lived side-by-side with the first farmers [67]. In Scandinavia, archaeological data shows a situation where foragers from the Pitted Ware Culture context existed in parallel with Neolithic farmers from the Funnel Beaker culture context [6].

## 3. Demographic changes during the late Neolithic and Bronze Age

The population turnover during the early Neolithic was not the last episode where a migrating group had a massive impact on peoples of Europe. The late Neolithic and Bronze Age were also times of large-scale migrations. Movements from the east have been revealed based on genomic data, first, unexpectedly, by the sequencing of the genome of a 24,000-year-old Siberian boy [“MA-1”, 74]. MA-1 showed affinities to modern-day Europeans and Native Americans but not to East Asians. A plausible scenario to explain this result was an ancient north Eurasian population which had contributed to both the first Americans and Europeans. The MA-1 individual belonged to a population that is lost today, but that likely populated much of northeastern Eurasia until some millennia ago. This group has likely contributed genetic material to Europe for a long time; for instance, more than 7,000 years ago, Scandinavian hunter-gatherers show affinities to MA-1 [22, 24, 7], but the impact on central and western Europe occurs in the late Neolithic [7, 8]. The Yamnaya herders from the Pontic-Caspian steppe have been suggested to have migrated to central Europe about 4,500 years ago, which likely brought that eastern genetic component to Europe [7, 8]. The steppe herders were themselves descendants from different hunter-gatherer groups from (modern-day) Russia [7] and the Caucasus [25]. This migration of the Yamnaya herders has been suggested to result in the rise of the late Neolithic Corded Ware culture in Central Europe and the spread of Indo-European languages [8, 7].

The late Neolithic and the Bronze Age were generally very dynamic times, which did not just involve the Corded Ware groups but also people associated with the Bell Beaker culture and the later Unetice culture as well as several other groups. The dynamics of these groups eventually spread the genetic material of the MA-1 descendants over most of western and northern Europe reaching the Atlantic coast and the British Isles [8, 7, 53] (Figure 2). these massive migrations homogenized European populations (Figure 1D) and reduced genetic differentiation similar levels as in modern Europeans [57]. In addition to the hunter-gatherers’ recolonization of Europe after the LGM, and the migrations connected with the Neolithic transition, the late Neolithic/Bronze Age migrations from the east are likely the third most influential event for the composition and gradients of genomic variation among modern-day Europeans (Figure 2).

## 4. Shaping European population structure after the Bronze Age

After the turnovers during the early Neolithic and Bronze Age periods, the genetic composition of populations in particular parts of Europe was already starting to become similar to the modern-day groups of the same regions (Figure 1D). This observation does not negate later migrations, but it shows that the populations involved were not as highly differentiated as during the Neolithic [57], when populations were almost as different from each other as modern-day continental groups [22]. Population history during the last three thousand years has been successfully studied using both ancient and modern DNA [e.g. 77, 78, 79, 80, 81, 82, 83]. Several studies have focused on the population history of the British Isles revealing population events during different time periods including the Iron Age [82, 83], the Roman period [80, 82], the Anglo-Saxon period [80, 83] and the viking migrations [80]. The populations of the European mainland were shaped by several small and large scale migrations [78, 79, 81], in addition to isolation-by-distance processes. Sources of these movements were both within and outside of Europe. Associating admixture events with particular historical events can be difficult, but effects of the “migration period” (during the first millennium CE; also known as the “Völkerwanderung”) have been suggested based on modern-day population-genetic data [81]. All these migration and admixture events produced the isolation-by-distance pattern that we see in modern Europeans [1, 2, 3]. It is also likely that the growing population size in Europe made later migrations less influential on demography since the relative fraction of migrants was decreasing.

## 5. Future directions for the genomic research of European prehistory

The field of ancient genomics is largely dependent on the availability of samples and DNA preservation; groups from more than 10,000 years ago and from non-temperate climates are under-represented among the published ancient genomes. Denser sampling – both on a geographic and a temporal scale – and better quality genomic data (e.g. high-coverage genomes) will help getting a deeper understanding of population structure during each period, identifying the exact source region of population movements (e.g. for the Neolithic migration) as well as allow for a detailed study of the mode of migrations (e.g. single wave versus multiple waves, sex-bias in admixture events). Methodological and technological advances – for working with both ancient and modern-day DNA – are going to reveal more insights into the processes that made the patterns of genomic variation across Europe what it is today. Although many open questions remain, the last few years of population genomic research have contributed in an extraordinary way to re-writing European history.

## Acknowledgments

We thank Amy Goldberg, Anders Götherström, Gülçsah Merve Dal Kılınç, Jan Apel, Jan Storå and Mehmet Somel for discussions and comments on earlier drafts of the review. This work was supported by an ERC starting grant (no. 311413) and a Swedish Research Council grant to M.J. Computations were performed on resources provided by SNIC through Uppsala Multidisciplinary Center for Advanced Computational Science (UPPMAX) under Project b2013203 [84].

*•* of special interest *••* of outstanding interest

